# DeepLncLoc: a deep learning framework for long non-coding RNA subcellular localization prediction based on subsequence embedding

**DOI:** 10.1101/2021.03.13.435245

**Authors:** Min Zeng, Yifan Wu, Chengqian Lu, Fuhao Zhang, Fang-Xiang Wu, Min Li

## Abstract

**Motivation:** Long non-coding RNAs (lncRNAs) are a class of RNA molecules with more than 200 nucleotides. A growing amount of evidence reveals that subcellular localization of lncRNAs can provide valuable insights into their biological functions. Existing computational methods for predicting lncRNA subcellular localization use k-mer features to encode lncRNA sequences. However, the sequence order information is lost by using only k-mer features.

**Results:** We proposed a deep learning framework, DeepLncLoc, to predict lncRNA subcellular localization. In DeepLncLoc, we introduced a new subsequence embedding method that keeps the order information of lncRNA sequences. The subsequence embedding method first divides a sequence into some consecutive subsequences, and then extracts the patterns of each subsequence, last combines these patterns to obtain a complete representation of the lncRNA sequence. After that, a text convolutional neural network is employed to learn high-level features and perform the prediction task. Compared to traditional machine learning models with k-mer features and existing predictors, DeepLncLoc achieved better performance, which shows that DeepLncLoc could effectively predict lncRNA subcellular localization. Our study not only presented a novel computational model for predicting lncRNA subcellular localization but also provided a new subsequence embedding method which is expected to be applied in other sequence-based prediction tasks.

**Availability:** The DeepLncLoc web server, source code and datasets are freely available at http://bioinformatics.csu.edu.cn/DeepLncLoc/, and https://github.com/CSUBioGroup/DeepLncLoc.

## 1 Introduction

Long non-coding RNAs (lncRNAs) are a type of large RNA molecules (more than 200 nucleotides) that are transcribed from DNA but not translated into proteins (Consortium, 2007; Lu, et al., 2018). LncRNAs play an important role in various biological processes including regulation of gene expression, alternative splicing, nuclear organization, and genomic imprinting (Moran, et al., 2012). For example, lncRNAs can bind to DNAs, RNAs, and proteins, and then perform their functions through these interactions (Esteller, 2011). LncRNAs can act as “miRNA sponge” to regulate the level of miRNA and then affect the expression of miRNA’s target (DiStefano, 2018). LncRNAs can regulate transcriptional activity or pathways under specific stimulation (Wang and Chang, 2011). Due to the complexity of molecular functions, lncRNAs-related studies have received a lot of attention (Lu, et al., 2019).

A growing amount of evidence reveals that the subcellular localization of lncRNAs can provide valuable insights into their functions (Carlevaro-Fita and Johnson, 2019). For example, lncRNA “XIST”, which locates in nucleus, interacts with the nuclear-matrix factor hnRN- PU and modulates nuclear architecture and trans-chromosomal interactions (Hacisuleyman, et al., 2014). LncRNA “lincRNA-p21”, which locates in cytoplasm, regulates JUNB and CTNNB1 translation in HeLa cells (Yoon, et al., 2012). LncRNA “ZFAS1”, which locates in ribosome, regulates mRNAs encoding proteins from the ribosomal complex (Hansji, et al., 2016). Thus, identification of lncRNA subcellular localizations is very important to understand lncRNA functions (Voit, et al., 2015).

Recently, some large databases of RNA-associated subcellular localization were released. Zhang et al. published a database, RNALocate (Zhang, et al., 2016), to collect the subcellular localization of different kinds of RNAs, which contains more than 23,100 RNAs with 42 subcellular localizations in 65 species. Mas Ponte et al. developed a database called LncATLAS to display the subcellular localization of lncRNAs (Mas-Ponte, et al., 2017). Wen et al. created a lncRNA subcellular localization database called lncSLdb (Wen, et al., 2018), which collects 14,973 subcellular localization information of lncRNAs from 3 species (human, mouse, and fruitfly).

However, only a few computational predictors for lncRNA subcellular localization have been proposed. To our knowledge, the first predictor is lncLocator (Cao, et al., 2018). LncLocator uses 4-mer features and high-level features extracted by stacked autoencoder, and feeds the two kinds of features into two kinds of classifiers (support vector machine and random forest), respectively. Then lncLocator uses an ensemble strategy to combine the results of different classifiers and get the final prediction. In their training process, lncLocator utilizes a supervised over-sampling algorithm to balance the ratio of different classes. The second predictor is iLoc-lncRNA (Su, et al., 2018). iLoc-lncRNA uses 8- mer features to encode lncRNA sequences. Considering the dimension of 8-mer features is too large, iLoc-lncRNA applies a feature selection method based on binomial distribution to select the most optimal features. Then iLoc-lncRNA feeds the most optimal features into support vector machine (SVM) to get the prediction results. The third predictor is DeepLncRNA (Gudenas and Wang, 2018). DeepLncRNA uses 2, 3, 4, and 5-mer features to encode lncRNA sequences, and adds additional features (RNA-binding motifs and genomic loci). Then the combined features are feed into a neural network to obtain the final prediction. Although these computational predictors achieve decent performance, several improvements can still be made. Encoding raw lncRNA sequences into discriminative features is very important in developing machine learning models. The flaw of these predictors is the use of only k-mer features to encode raw lncRNA sequences. Apparently, using only k-mer features cannot keep the sequence order information of the raw lncRNA sequence.

To overcome the limitation, we developed DeepLncLoc, a new deep learning-based predictor for subcellular localization of lncRNAs. In the predictor, we proposed a new feature embedding method that keeps the order information of lncRNA sequences (see “Section 2.3” for details). The main idea of the new feature embedding method is encoding a complete RNA sequence by using the combination of its subsequence embedding. In DeepLncLoc, we divided a sequence into some consecutive subsequences, and then extracted the patterns of each subsequence by using an average pooling layer; last combined these patterns to obtain a complete representation of the lncRNA sequence. After obtaining the complete representation, a text convolutional neural network (textCNN) was applied to learn high-level features and perform the prediction task. Different from traditional machine learning models with k-mer features in previous studies, DeepLncLoc has two advantages, i) by using the new subsequence embedding method, the input lncRNA sequence keeps the sequence order information, ii) textCNN has a more powerful capability of high-level feature extraction.

We conducted extensive experiments to evaluate the performance of DeepLncLoc. Comparison with traditional machine learning models with different k-mer features demonstrated the advantages of using subsequence embedding to encode the whole lncRNA sequence instead of using only k-mer features. Comparison with existing predictors on an independent test set showed the capability of DeepLncLoc to predict subcellular localization of lncRNAs. Moreover, we investigated the effects of different species. Finally, we developed a user-friendly web server.

## 2 Methods

### 2.1 Dataset

Similar to previous studies, we retrieved known subcellular localization information of lncRNA from RNALocate database (Zhang, et al., 2016). The current version of RNALocate collects 42,190 manually curated RNA-associated subcellular localization entries with experimental evidence. It contains more than 23,100 RNAs with 42 subcellular localizations in 65 species. We generated a benchmark dataset to train and test our model by the following procedure:

1. All 42,190 manually curated RNA-associated subcellular localization entries are downloaded from RNAlocate database;
2. Total 2,383 manually curated lncRNA-associated subcellular localization entries are selected from 42,190 manually curated RNA-associated subcellular localization entries;
3. Some lncRNAs have multiple entries in the extracted entries, we merged these entries with the same gene name. Then we removed the lncRNAs that do not have sequence information in NCBI and Ensembl.
4. Because most lncRNAs only have one subcellular localization, we selected the lncRNAs that are located in one location for model construction in the study.
5. The filtered dataset covers seven different subcellular localizations. Two of seven subcellular localizations only have a very small number of samples (less than 10). Thus we removed these lncRNAs that are located in the two subcellular localizations.

Finally, we constructed a benchmark dataset of 857 lncRNAs, covering 5 subcellular localizations including nucleus, cytosol, ribosome, cytoplasm, and exosome (see Supplementary Fig. S1). Table 1 lists the distribution of the constructed benchmark dataset.

**Table 1.** Distribution of the constructed benchmark dataset.

| Subcellular localization | # of samples |
| --- | --- |
| Cytoplasm | 328 |
| Nucleus | 325 |
| Ribosome | 88 |
| Cytosol | 88 |
| Exosome | 28 |
| Total | 857 |

### 2.2 Limitations of using only k-mer features to encode RNA sequences

Before putting raw RNA sequences into a machine learning or deep learning model, RNA sequences need to be encoded as numeric vectors. There are two kinds of widely used RNA sequence embedding methods. The first one is encoding each nucleotide into a 4-dimensional one-hot vector. The A, C, G and U are encoded with a one-hot vector of (1, 0, 0, 0), (0, 1, 0, 0), (0, 0, 1, 0) and (0, 0, 0, 1), respectively (Pan, et al., 2019). Then the four types of vectors are used to encode RNA sequences. However, using one-hot encoding has two disadvantages in practice. The first disadvantage is that one-hot vector is sparse, i.e., only a small fraction of features contributes to the prediction task. The second disadvantage is that using one-hot encoding is difficult to accurately represent the similarity between different nucleotides. The second method is using k-mer features to encode RNA sequences. The k-mer feature encoding method is very simple to implement, and it maps lncRNA sequences with variable-length to a vector with a fixed dimension. Thus, k-mer feature encoding method is the most widely used method in the prediction of lncRNA subcellular localization. Previous methods (LncLocator (Cao, et al., 2018), iLoc-lncRNA (Su, et al., 2018) and DeepLncRNA (Gudenas and Wang, 2018)) use k-mer features for lncRNA embedding. Formally, we assume a lncRNA sequence is represented as:

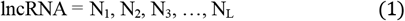

where L denotes the length of the lncRNA, N_i_ is one of the four nucleotide bases (A, C, G and U) in the *i* position of the lncRNA sequence.

For a given k, k-mer features represent the frequency of individual k-mer from lncRNA sequences. We take 3-mer as an example, each position can take four nucleotide bases (A, C, G and U), thus we have 4^3^, i.e., 64 3-mer features (AAA, AAC, …, UUU). Then we can use a 64-dimensional vector to represent a lncRNA sequence, and each dimension is used to record the occurrence time of a certain 3-mer. Fig. 1 plots the k-mer encoding method for a single RNA sequence. The k-mer feature encoding method is very simple to understand and implement. But there is a disadvantage of using k-mer features. Namely, k-mer feature encoding method lost order information of the raw lncRNA sequence. K-mer features encoding method is only concerned with the occurrence of the k-mer and ignores the position of k-mer in the raw lncRNA sequence. For example, RNA A is “ACACACGCGC”, 3-mer features of RNA A are {ACA, CAC, ACA, CAC, ACG, CGC, GCG, CGC}; we reverse the RNA sequence to obtain RNA B “CGCGCACACA”, the 3-mer features of RNA B are {CGC, GCG, CGC, GCA, CAC, ACA, CAC, ACA}. It can be seen that the order of the two RNA sequences is reversed, but their 3-mer features are very similar. The difference between the two 3- mer features is only one 3-mer (“ACG” in RNA A versus “GCA” in RNA B). When using the 64-dimensional 3-mer vector to encode the two lncRNA sequences, only two dimensions are different.

**Fig. 1.**
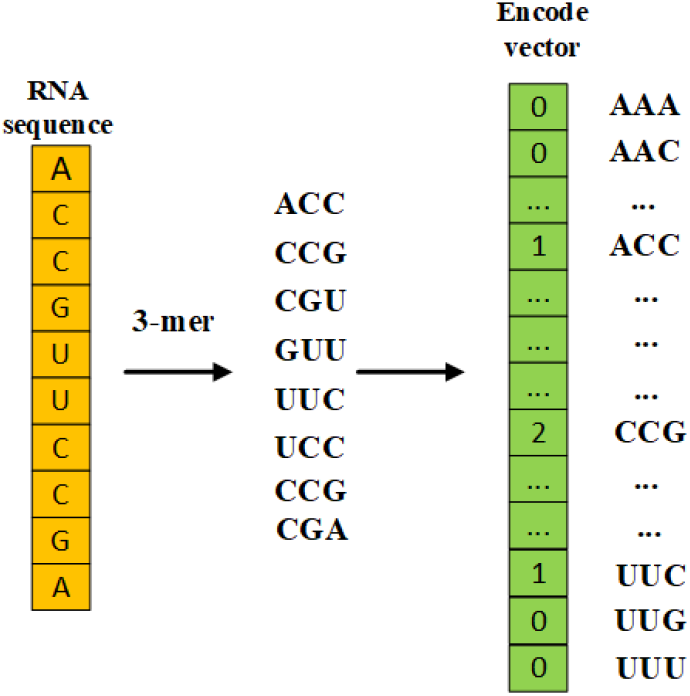
Illustration of the k-mer encoding method for single RNA sequence, where k is set to 3. The example RNA sequence is “ACCGUUCCGA”.

### 2.3. Subsequence embedding

In order to tackle the limitation, we proposed an effective subsequence embedding method to keep the sequence order information of lncRNAs. The main idea is that we split a lncRNA sequence into some consecutive subsequences with no overlap between the subsequences, and then extract the patterns of each subsequence; last we combine these patterns to obtain a complete representation of the lncRNA sequence. In this way, we can keep the sequence order information. The idea is motivated by spatial pyramid pooling-net (He, et al., 2015), He et al. proposed spatial pyramid pooling-net to obtain the features from arbitrary sub-images to generate fixed-length representations for the entire image. We transferred and modified their idea to encode lncRNA sequences.

We split a lncRNA sequence into *m* consecutive subsequences, thus we denote a lncRNA sequence:

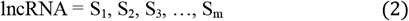

where Si is the *i*th subsequence. We denote Lsi is the length of Si. After dividing a lncRNA sequence into *m* subsequences, the next step is encoding these subsequences. Word embedding techniques have shown promise in many natural language processing applications including text classification, sentiment analysis, and part-of-speech tagging. We used a word embedding technique to encode subsequences. Specifically, we pre-trained lncRNA sequences in our dataset to obtain the distribution representation of k-mer by using word2vec technique, and then used the distribution representation of k-mer features to represent subsequences. Word2vec is a popular word embedding technique (Mikolov, et al., 2013). It aims at learning a dense vector automatically for each word in a corpus. The word2vec technique has two models: skip-gram and continuous bag of words (CBOW) model. The skip-gram model uses the central word to predict context words. In the training process, we maximized the co-occurrence likelihood function of the central word and corresponding context words. In our study, we used gensim library to learn k-mer features of lncRNA sequences (Rehurek and Sojka, 2010). The parameter k is chosen from {1, 2, 3, 4, 5, 6} to find the best value.

The steps of subsequence embedding (see the subsequence embedding part in Fig. 2) are described as follows:

1. We used gensim library to learn representation vectors of k-mer (D-dimension) of all lncRNA sequences in our database.
2. For a given lncRNA, we split it into *m* subsequences, the length of each subsequence is L_si_.
3. According to the k value in step 1, for each subsequence, used k-mer features to encode it.
4. Found the pre-trained vector of each k-mer, and then combined these vectors into a matrix as the representation of a subsequence.

**Fig. 2.**
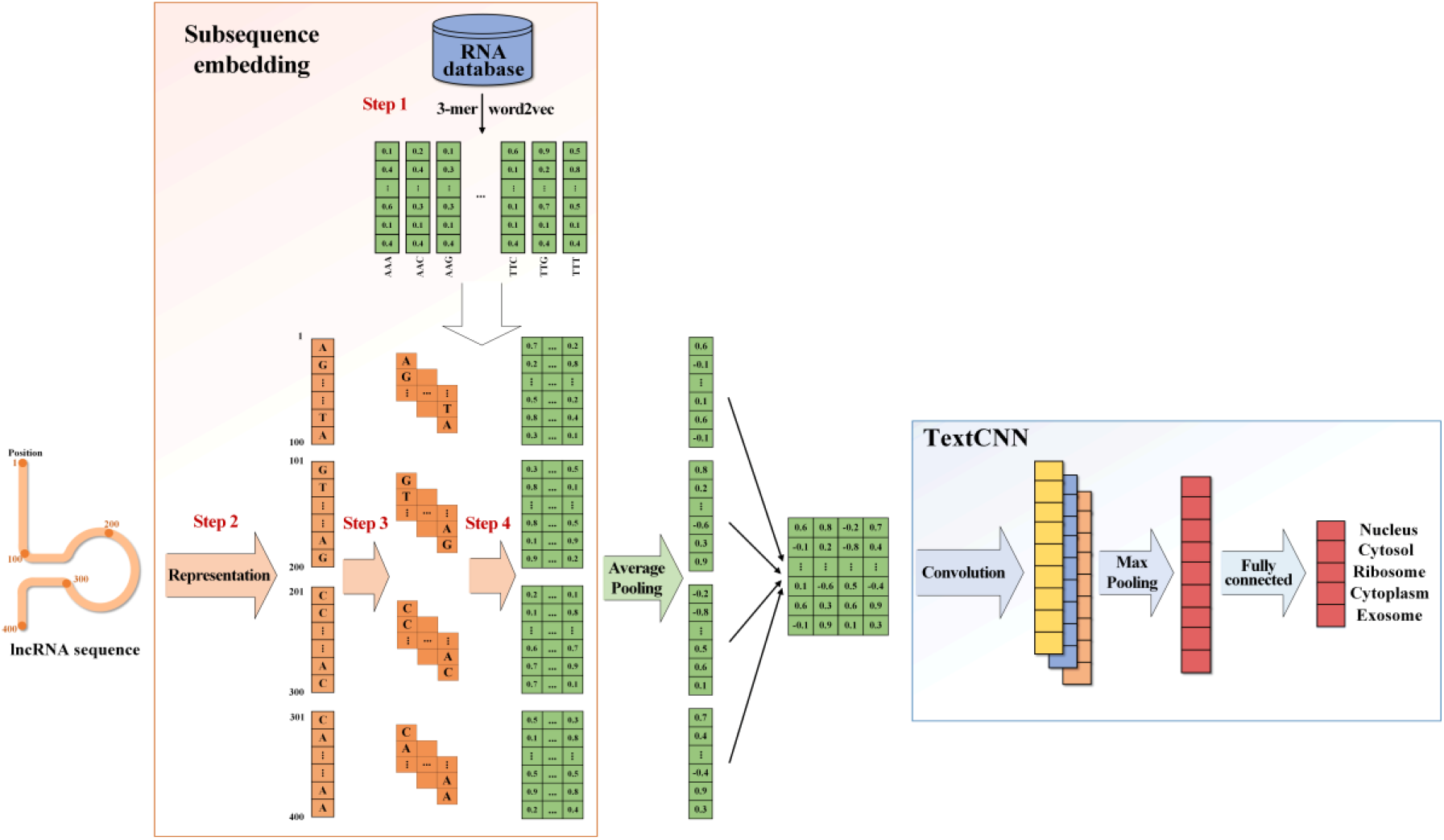
Illustration of the deep neural network structure. This figure is only an example. The network structure consists of three parts: subsequence embedding, an average pooling layer, and a textCNN. The input is a lncRNA sequence with a length of 400. The lncRNA sequence is split into 4 subsequences. The sequence embedding part has four steps. After subsequence embedding, we use an average pooling layer to extract the patterns of each subsequence. Then we combine these patterns together to obtain a matrix as the representation of the whole lncRNA sequence. Last, a textCNN is employed to learn high-level features and perform the prediction task.

Last, we converted each lncRNA subsequence into a matrix whose dimension is (L_si_-2) * D (L_si_ is the length of each subsequence), which is the actual input for our deep learning model.

### 2.4. Network architecture

So far we have obtained the representation of each subsequence. The question then arises: how can we predict the subcellular localization by using the representation of subsequences. We have *m* subsequences, and the representation of each subsequence is a matrix whose dimension is D * (L_si_-2). If we put them together directly, the dimension is N * D * (Lsi - 2), which has two disadvantages. First, the length of different subsequence Lsi in different lncRNA sequence is not the same. If we put them together directly, we must pad them to the same length. It means we have to fill a lot of zeros at the end of the raw sequence, which brings many meaningless between subsequences and vectors with all zeros. Second, the dimension is too large after putting them together directly, which causes a lot of computational waste. To tackle the two limitations, we use an average pooling layer to extract the patterns in each channel of the subsequence. By using the average pooling layer, the dimension of each sequence is reduced from D * (L_si_-2) to D. It can be seen that D is the dimension of the pre-trained vector of k-mer, and has no relationship with the length of lncRNA subsequence L_si_. By using this method, we do not need to pad with zeros and reduce the dimensionality.

After obtaining the representation of each subsequence by using an average pooling layer, we combined them together to obtain the complete representation of the whole lncRNA sequence. Then the next step is predicting the subcellular localization. TextCNN is a kind of powerful deep learning network structure that is used for text classification. Traditional CNNs are two-dimensional CNNs that are used to process twodimensional image data. Actually, a text can be treated as a onedimensional image, so that we can use one-dimensional CNN to extract features of the text. TextCNN uses a one-dimensional convolutional layer and a max-pooling layer to extract features of sequence (Kim, 2014). Inspired by its success in bioinformatics (Zeng, et al., 2019), we used textCNN to extract features of the complete representation. Specifically, we have *m* subsequences, and the representation of each subsequence is D. We combined them together to form a matrix whose dimension is N*D to represent the whole sequence. The representation of the lncRNA sequence can be treated as a one-dimensional image, the width is N, the height is 1, and the channel is D. To extract high-level features, textCNN uses three convolutional kernels (sizes=1, 3, 5) to capture the correlation of adjacent nucleotides. Then textCNN performs a maxpooling layer on all channels to obtain the most remarkable features and reduce the dimension of the output vector. Last, the output vectors of the max-pooling layer are concatenated together as the input of a fully connected layer with a softmax function to perform the final prediction. Fig. 2 gives a schematic view of the whole network structure.

### 2.5 Evaluation metrics

Similar to previous studies (Cao, et al., 2018; Gudenas and Wang, 2018; Su, et al., 2018), we used accuracy (ACC), Macro F-measure, and area under the receiver operator characteristic curve (AUC) as evaluation metrics to evaluate DeepLncLoc and other methods in the study.

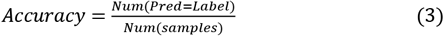

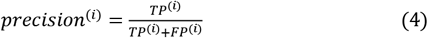

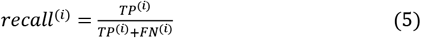

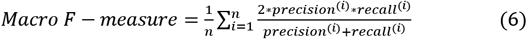

where TP^(i)^, FP^(i)^, and FN^(i)^ represent the number of true positives, false positives, and false negatives of the class *i*, respectively.

### 2.6 Implementation details

DeepLncLoc is implemented with PyTorch (Paszke, et al., 2017). The loss function used in DeepLncLoc is the focal loss of non-α-balanced form (Lin, et al., 2017). It is used for object detection to address this class imbalance problem. It is defined as follows:

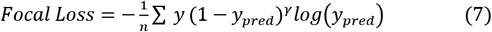

where n is the number of training samples, y is the true label, and ypred is the predicted label, *γ* is the focusing parameter (we set *γ* to 2, according to Lin’s paper).

Skip-gram model (Mikolov, et al., 2013) is used to pre-train the vectors of k-mer for embedding. In textCNN, three convolutional kernels (sizes=1, 3, 5, filter number=128) are used to extract the high-level features of adjacent nucleotides. The fully connected layer in the classification part has 384 neurons. To avoid overfitting, dropout rates of 0.3 and 0.5 are applied in the embedding layer and the fully connected layer, respectively. Finally, we trained DeepLncLoc using the Adaptive Momentum optimizer, the initial learning rate is set to 0.001.

## 3 Results

### 3.1 Hyper-parameter optimization for DeepLncLoc

We used 5-fold cross-validation (5-fold CV) to tune the hyperparameters of DeepLncLoc based on the value of Macro F-measure. In our model, many hyper-parameters affect the computational results, such as the parameter *k*, the number of subsequences, the dimension of the pre-trained vector of k-mer, initial learning rate, and kernel sizes. In the study, we cared about most is the effect of subsequence embedding on computational results. Thus, we considered the parameter *k*, the number of subsequences *m* and the dimension of the pre-trained vector of k-mer *d* as the major tuning hyper-parameters. A grid search strategy is applied to find the best combination of the three hyper-parameters. The parameter *k* was chosen from {1, 2, 3, 4, 5, 6}, the number of subsequences *m* was chosen from {16, 32, 64, 128, 256} and the dimension of pre-trained vector *d* was chosen from {64, 128}. We tuned these hyper-parameters to find the final model parameters (see Supplementary Table S1). From Table S1, it is very hard to determine the parameters directly. We analyzed and found that the performance is unstable when *k* and *m* are too high or too low. In order to ensure the generalization of DeepLncloc, *k, m*, and *d* are set to 3, 64, and 64, respectively. In this setting, the ACC, Macro F-measure and AUC obtained by DeepLncLoc are 0.548, 0.421 and 0.820, respectively.

### 3.2 Comparison with traditional machine learning classifiers with different k-mer features

Considering that traditional machine learning classifiers with k-mer features are widely used in the prediction of lncRNA subcellular localization, we compared DeepLncLoc with three traditional machine learning models including SVM, random forest (RF) and logistic regression (LR). The parameter k in these machine learning models was chosen from {3, 4, 5, 6}. We did not consider the lower and higher k because much lower or higher k will increase the risk of underfitting or overfitting. For example, the dimension of 2-mer features is 4^2^, i.e., 16, which hardly encodes the diversity of all sequences in the database. In this case, the model has a high risk of underfitting. The dimension of 7-mer fea-tures is 4^7^, i.e., 16,384, which is far beyond the number of all samples. In this case, the model has a high risk of overfitting. The results are shown in Table 2.

**Table 2.** Performance of DeepLncLoc and different machine learning models with different k-mer features.

|  | Model | ACC | Macro F-measure | AUC |
| --- | --- | --- | --- | --- |
| $k=3$ | SVM | 0.481 | 0.224 | 0.794 |
|  | RF | 0.480 | 0.305 | 0.777 |
|  | LR | 0.497 | 0.267 | 0.813 |
| $k=4$ | SVM | 0.486 | 0.223 | 0.808 |
|  | RF | 0.508 | 0.327 | 0.788 |
|  | LR | 0.469 | 0.289 | 0.775 |
| $k=5$ | SVM | 0.499 | 0.271 | 0.811 |
|  | RF | 0.497 | 0.282 | 0.786 |
|  | LR | 0.446 | 0.290 | 0.728 |
| $k=6$ | SVM | 0.496 | 0.245 | 0.809 |
|  | RF | 0.483 | 0.280 | 0.772 |
|  | LR | 0.479 | 0.335 | 0.767 |
|  | DeepLncLoc | **0.548** | **0.421** | **0.820** |

Note: The best performance values are highlighted in bold.

From Table 2, first noted that the performance of each machine learning model with different k-mer features is different. We can see that the best performance of SVM, RF is achieved when k=5, 4, respectively. For LR, the highest ACC, Macro F-measure, AUC are achieved when k=3, 6, 3, respectively. Second, all evaluation metrics obtained by DeepLncLoc are higher than other machine learning classifiers. The ACC and Macro F-measure of DeepLncLoc are significantly higher than the other ma-chine learning methods. The AUC of DeepLncLoc is slightly higher than the other machine learning methods. Fig. 3 plots the ROC curves of DeepLncLoc and other machine learning methods with the highest AUC. It is obvious that DeepLncLoc has the highest AUC value on each class. This indicated that our proposed computational approach is better than traditional machine learning models with k-mer features.

**Fig. 3.**
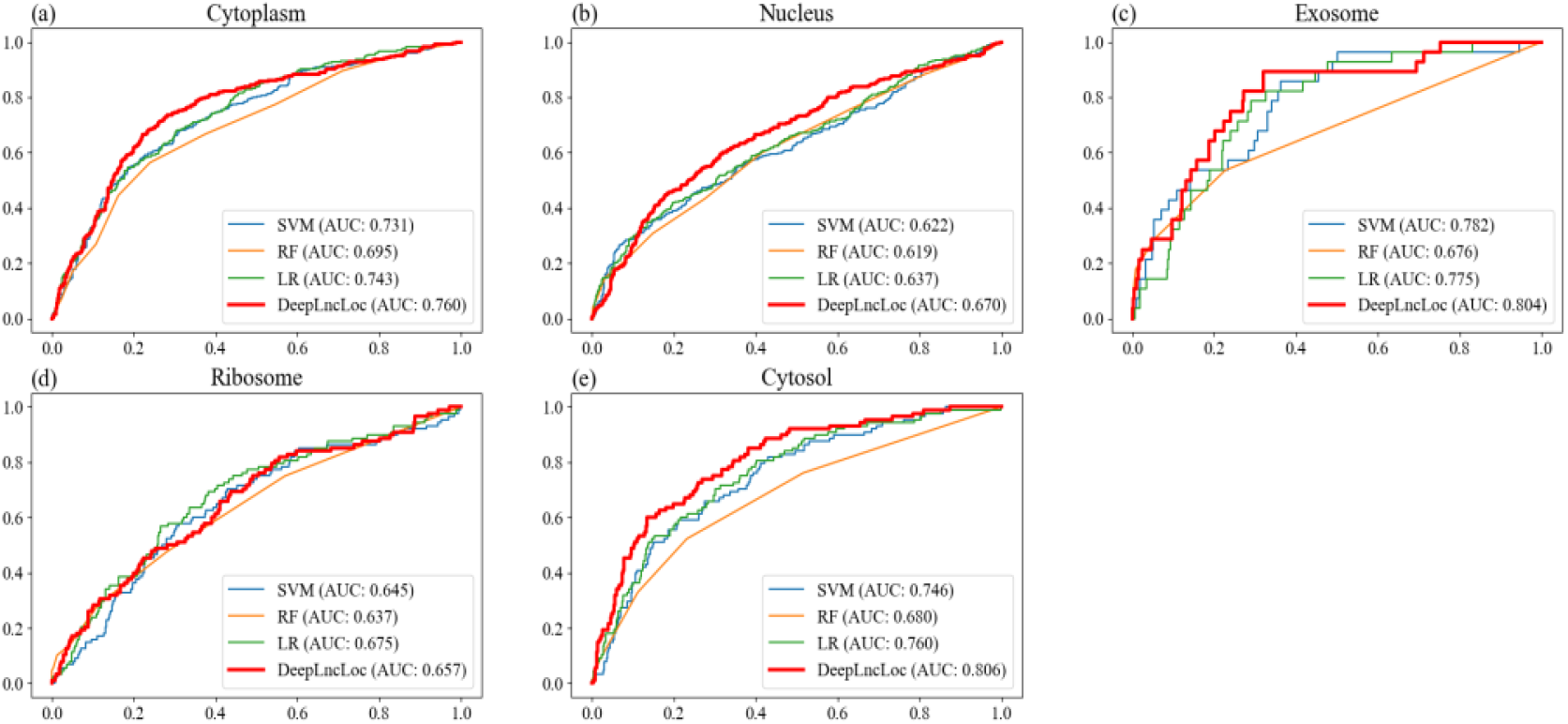
The ROC curves of DeepLncLoc, SVM (k=5), RF (k=4), and LR (k=3) for each class. (a) Cytoplasm, (b) Nucleus, (c) Exosome, (d) Ribosome, (e) Cytosol.

### 3.3 Comparison with current predictors

The 5-fold CV is applied in our previous experiments. To further evaluate the performance of DeepLncLoc in predicting the subcellular localization of lncRNAs, we compared DeepLncLoc with current predictors by using a stand-alone test set.

We selected current predictors follow these criteria: i) availability of web server or stand-alone version; ii) input that only needs lncRNA sequences; and iii) outputs that include predictive scores for subcellular localization. Consequently, lncLocator (Cao, et al., 2018) and iLoc-lncRNA (Su, et al., 2018) satisfy these criteria. LncLocator can predict 5 subcellular localizations of lncRNAs, including nucleus, cytoplasm, cytosol, ribosome, and exosome. iLoc-lncRNA can predict 4 subcellular localizations of lncRNAs, including nucleus, cytoplasm, ribosome, and exosome. We used the web server of lncLocator (available at http://www.csbio.sjtu.edu.cn/bioinf/lncLocator/) and iLoc-lncRNA (available at http://lin-group.cn/server/iLoc-LncRNA/download.php) for comparison.

We compared DeepLncLoc with the two predictors (lncLocator and iLoc-lncRNA) by using an independent test set. The test set was created from another lncRNA subcellular localization database lncSLdb and recent literature. Since lncSLdb database only collects 5 subcellular localizations: nucleus, chromosome, cytoplasm, nucleoplasm, and ribosome, and does not have records in the subcellular localization of cytosol and exosome. Thus, we randomly selected some samples from 3 subcellular localizations (nucleus, cytoplasm, and ribosome) in lncSLdb database. To obtain other samples from the subcellular localization of cytosol and exosome, we searched some recent literature in the PubMed database using the following keywords: lncRNA and each subcellular localization, and then obtained lncRNA sequences from NCBI database. we used the cd-hit tool to remove the redundant sequences with a cutoff of 90%. Last, the test set contains 20 samples from cytoplasm, 20 samples from nucleus, 10 samples from ribosome, 10 samples from cytosol, and 7 samples from exosome (see Supplementary Table S2). All lncRNA sequences in the independent test set are not used for the construction of DeepLncLoc. The independent test set can be accessed at https://github.com/CSUBioGroup/DeepLncLoc/tree/master/Independent_test_set.

**Table 3.**
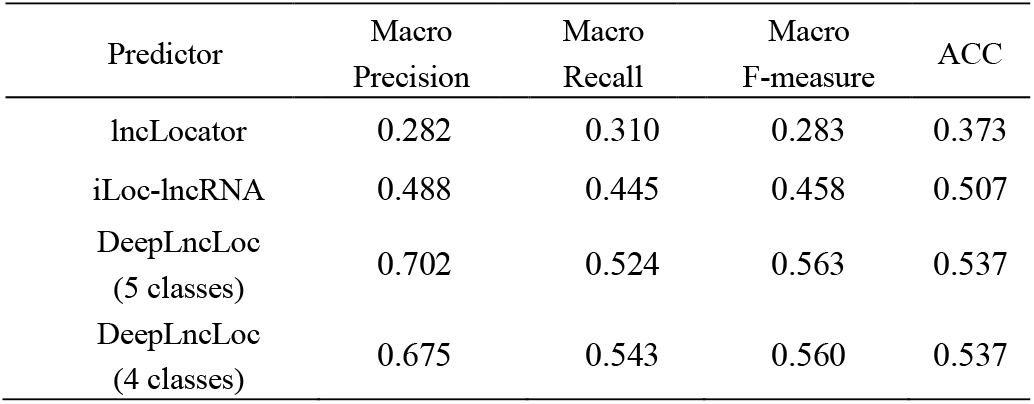
Comparison of the prediction performance of DeepLncLoc with lncLocator and iLoc-lncRNA on the test set.

The confusion matrices of DeepLncLoc and lncLocator are shown in Supplementary Fig. S2. Since iLoc-lncRNA treats cytoplasm and cytosol as one category, it only predict four classes (nucleus, cytoplasm, ribosome, and exosome). To make the comparison fair, we treated cytoplasm and cytosol as one category when we compared DeepLncLoc with iLoc-lncRNA. The confusion matrices of DeepLncLoc and iLoc-lncRNA are shown in Supplementary Fig. S3. In Supplementary Figs. S2 and S3, each row represents the true class while each column represents the predicted class. The diagonal elements represent the number of samples that are predicted correctly. Out of the 68 lncRNAs, our method predicted correct subcellular localization for 36 of them, which is far more accurate than lncLocator (25) and slightly higher than iLoc-lncRNA (34). The results of DeepLncLoc, lncLocator, and iLoc-lncRNA are reported in Table 3. Clearly, the accuracy of DeepLncLoc is higher than lncLocator and iLoc-lncRNA. The Macro Precision, Macro Recall, Macro F-measure of DeepLncLoc (5 classes) are 0.702, 0.524, and 0.563, respectively, which are significantly higher than those of lncLocator (0.282, 0.310, and 0.283). Similar results are observed when we compared DeepLncLoc (4 classes) with iLoc-lncRNA. All results suggested that the DeepLncloc may serve as a useful tool to predict the subcellular localization of lncRNAs. We gave the detailed prediction results of DeepLncLoc, lncLocator, and iLoc-lncRNA on the test set (see Supplementary Table S3). Precision, recall, F-measure of DeepLncLoc, lncLocator, and iLoc-lncRNA for each class on the test set are reported in Tables 4 and 5. We observed that the F-measures of DeepLncLoc for nucleus, ribosome, cytosol, and exosome are higher than those of lncLocator, and the F-measures of DeepLncLoc for cytoplasm is lower than that of lncLocator. This phenomenon has been observed when we compared DeepLncLoc with iLoc-lncRNA. In addition, we also noted that none of samples in exosome have been correctly recognized by lncLocator, which lead to very bad prediction results for exosome. A possible explanation is that there are too many samples of cytoplasm in the training set of lncLocator and iLoc-lncRNA. The machine learning model will naturally give more preference to cytoplasm, resulting in a bad performance for the other classes. Thus lncLocator and iLoc-lncRNA tend to classify other subcellular localization to cytoplasm.

**Table 4.** Precision, recall, F-measure of DeepLncLoc and lncLocator for each class on the test set.

| Predictor | lncLocator |  |  | DeepLncLoc |  |  |
| --- | --- | --- | --- | --- | --- | --- |
|  | Precision | Recall | F<sub>1</sub> | Precision | Recall | F<sub>1</sub> |
| Cytoplasm | 0.484 | 0.750 | 0.588 | 0.778 | 0.350 | 0.483 |
| Nucleus | 0.308 | 0.200 | 0.242 | 0.400 | 0.800 | 0.533 |
| Ribosome | 0.333 | 0.200 | 0.250 | 0.500 | 0.400 | 0.444 |
| Cytosol | 0.286 | 0.400 | 0.333 | 0.833 | 0.500 | 0.625 |
| Exosome | 0.000 | 0.000 | 0.000 | 1.000 | 0.571 | 0.727 |

Note: F<sub>1</sub> represents F-measure.

**Table 5.** Precision, recall, F-measure of DeepLncLoc and iLoc-lncRNA for each class on the test set.

| Predictor | iLoc-lncRNA |  |  | DeepLncLoc |  |  |
| --- | --- | --- | --- | --- | --- | --- |
|  | Precision | Recall | F<sub>1</sub> | Precision | Recall | F<sub>1</sub> |
| Cytoplasm | 0.553 | 0.700 | 0.618 | 0.800 | 0.400 | 0.533 |
| Nucleus | 0.467 | 0.350 | 0.400 | 0.400 | 0.800 | 0.533 |
| Ribosome | 0.333 | 0.300 | 0.316 | 0.500 | 0.400 | 0.444 |
| Exosome | 0.600 | 0.429 | 0.500 | 1.000 | 0.571 | 0.727 |

Note: F<sub>1</sub> represents F-measure.

### 3.4 The effects of different species

In addition, we investigated whether the species has an impact on classification results. The dataset covers six different species and the species distribution of lncRNAs is shown in Supplementary Table S5. Four species only have one or two lncRNAs, thus we only used two species (Homo sapiens and Mus musculus) for analysis. Homo sapiens group contains 461 lncRNAs and Mus musculus group contains 391 samples. Supplementary Fig. S4 plots the performance of DeepLncLoc on the two species. As shown in this figure, the ACC and AUC of Homo sapiens group are 0.547 and 0.823, respectively, which is slightly higher than those of Mus musculus group (0.503 and 0.774).

### 3.5 DeepLncLoc web server

A web server that implements DeepLncLoc is freely available at http://bioinformatics.csu.edu.cn/DeepLncLoc/. DeepLncLoc requires a lncRNA sequence with more than 200 and less than 100,000 nucleotides as input. Then click on the submit button to see the predicted results. The results have one table and one sentence, and will be shown on the screen of your computer. The table has five columns and each column represents the name of subcellular localization and corresponding probability. Last, the final predicted subcellular localization is marked red to show. Usually, DeepLncLoc takes less than 5 seconds to predict the subcellular localization of a lncRNA sequence.

## 4 Discussion and conclusion

Prediction of lncRNA subcellular localization can help to understand the complex biological functions of lncRNAs. However, all existing computational tools use k-mer features to encode lncRNA sequences, which lost the sequence order information. In this paper, we proposed DeepLncLoc, an open-sourced deep learning model, for predicting subcellular localization of lncRNAs. DeepLncLoc uses a novel subsequence embedding method to encode lncRNA sequences, then applied a textCNN to perform the classification task. Compare with previous studies, DeepLncLoc has two novel design ideas: i) it can keep the sequence order information of lncRNA sequences by using subsequence embedding; ii) textCNN can automatically capture high-level features from the combination of the patterns of all subsequences.

For the comparison of DeepLncLoc with other traditional machine learning methods, DeepLncLoc outperforms all traditional machine learning models with different k-mer features in terms of accuracy, Macro F-measure, and AUC. This implies that our proposed subsequence embedding method might be better than traditional k-mer features. Further comparison of DeepLncLoc with existing predictors by using an independent test set, DeepLncLoc outperforms existing predictors in terms of classification accuracy and Macro F-measure. This indicates that DeepLncLoc may serve as a useful tool to predict the subcellular localization of lncRNAs.

While our results are promising, several improvements can still be made. We would like to point out the following limitations of DeepLncLoc:

1. Because the majority of lncRNAs in RNALocate database only have one subcellular localization, thus we only chose the lncRNAs that only have one subcellular localization for training and testing in this study. However, in reality, many lncRNAs have multiple subcellular localizations. Therefore, in future work, if we can collect more labeled lncRNAs with multiple subcellular localizations, we can expand the dataset to train a more powerful model.
2. We only used lncRNA sequence-based features in our model for training and did not consider other biological information. There are some useful features that could be integrated for better predicting the subcellular localization (Zeng, et al., 2019; Zhang, et al., 2019). For example, Gudenas et al. used k-mer features, RNA-binding motifs and genomic loci to predict the subcellular localization of lncRNAs. Thus, in the future, we plan to incorporate other biological information to deep neural networks.
3. To reduce computational cost and runtime, we did not use a very complex deep learning model to extract features and perform the classification task. With the development of deep learning techniques, more and more powerful network architecture will be proposed. Therefore, using more powerful network structure to predict the subcellular localization is a promising future direction.
4. Classification for the minority class of subcellular localization (e.g. ribosome) is a challenging problem. This could be due to two reasons. First, there are too few samples in the minority class, it causes that our model cannot capture the patterns of the minority class. Second, the class distribution is imbalanced, the classifier tends to bias to the majority class (e.g. nucleus) and hence leads to a loss of predictive performance for the minority class (He and Garcia, 2008).

The variable-length of lncRNA sequences is hard to address in most existing computational methods. Even though our analysis was limited to predicting the subcellular localization of lncRNAs, we obtained promising results. We believe that the subsequence embedding method in DeepLncLoc can be used as a general representation method of RNA and DNA sequences. It is expected to be applied to other related variable-length sequence problems, such as prediction of mRNA subcellular localization (Yan, et al., 2019), prediction of DNA N4-methylcytosine sites (Wei, et al., 2018), RNA shape prediction (Mautner, et al., 2019).

## Funding

This work was supported in part by the National Natural Science Foundation of China under Grants (No. 61832019 and No. 61728211), the 111 Project (No. B18059), Hunan Provincial Science and Technology Program (2019CB1007).

*Conflict of Interest: none declared*

## Notes

https://github.com/CSUBioGroup/DeepLncLoc

### Competing Interest Statement

The authors have declared no competing interest.

## References

Cao, Z., et al. (2018) The lncLocator: a subcellular localization predictor for long non-coding RNAs based on a stacked ensemble classifier, Bioinformatics, 34, 2185–2194.

Carlevaro-Fita, J. and Johnson, R. (2019) Global positioning system: understanding long noncoding RNAs through subcellular localization, Molecular cell, 73, 869–883.

Consortium, E.P. (2007) Identification and analysis of functional elements in 1 % of the human genome by the ENCODE pilot project, Nature, 447, 799.

DiStefano, J.K. (2018) The emerging role of long noncoding RNAs in human disease. In, Disease Gene Identification. Springer, pp. 91–110.

Esteller, M. (2011) Non-coding RNAs in human disease, Nature Reviews Genetics, 12, 861.

Gudenas, B.L. and Wang, L. (2018) Prediction of lncRNA subcellular localization with deep learning from sequence features, Scientific reports, 8, 16385.

Hacisuleyman, E., et al. (2014) Topological organization of multichromosomal regions by the long intergenic noncoding RNA Firre, Nature structural & molecular biology, 21, 198.

Hansji, H., et al. (2016) ZFAS1: a long noncoding RNA associated with ribosomes in breast cancer cells, Biology direct, 11, 62.

He, H. and Garcia, E.A. (2008) Learning from imbalanced data, IEEE Transactions on Knowledge & Data Engineering, 1263–1284.

He, K., et al. (2015) Spatial pyramid pooling in deep convolutional networks for visual recognition, IEEE transactions on pattern analysis and machine intelligence, 37, 1904–1916.

Kim, Y. (2014) Convolutional neural networks for sentence classification, arXiv preprint arXiv:1408.5882.

Lin, T.-Y., et al. (2017) Focal loss for dense object detection. Proceedings of the IEEE international conference on computer vision. pp. 2980–2988.

Lu, C., et al. (2019) Predicting human lncRNA-disease associations based on geometric matrix completion, IEEE Journal of Biomedical and Health Informatics.

Lu, C., et al. (2018) Prediction of lncRNA-disease associations based on inductive matrix completion, Bioinformatics, 34, 3357–3364.

Mas-Ponte, D., et al. (2017) LncATLAS database for subcellular localization of long noncoding RNAs, Rna, 23, 1080–1087.

Mautner, S., et al. (2019) ShaKer: RNA SHAPE prediction using graph kernel, Bioinformatics, 35, i354–i359.

Mikolov, T., et al. (2013) Efficient estimation of word representations in vector space, arXiv preprint arXiv: 1301.3781.

Moran, V.A., Perera, R.J. and Khalil, A.M. (2012) Emerging functional and mechanistic paradigms of mammalian long non-coding RNAs, Nucleic acids research, 40, 6391–6400.

Paszke, A., et al. (2017) Automatic differentiation in pytorch.

Rehurek, R. and Sojka, P. (2010) Software framework for topic modelling with large corpora. In Proceedings of the LREC 2010 Workshop on New Challenges for NLP Frameworks. Citeseer.

Su, Z.-D., et al. (2018) iLoc-lncRNA: predict the subcellular location of lncRNAs by incorporating octamer composition into general PseKNC, Bioinformatics, 34, 4196–4204.

Voit, E.O., Martens, H.A. and Omholt, S.W. (2015) 150 years of the mass action law, PLoS computational biology, 11, e1004012.

Wang, K.C. and Chang, H.Y. (2011) Molecular mechanisms of long noncoding RNAs, Molecular cell, 43, 904–914.

Wei, L., et al. (2018) Exploring sequence-based features for the improved prediction of DNA N4-methylcytosine sites in multiple species, Bioinformatics, 35, 1326–1333.

Wen, X., et al. (2018) lncSLdb: a resource for long non-coding RNA subcellular localization, Database, 2018.

Yan, Z., Lécuyer, E. and Blanchette, M. (2019) Prediction of mRNA subcellular localization using deep recurrent neural networks, Bioinformatics, 35, i333–i342.

Yoon, J.-H., et al. (2012) LincRNA-p21 suppresses target mRNA translation, Molecular cell, 47, 648–655.

Zeng, M., et al. (2019) A deep learning framework for identifying essential proteins by integrating multiple types of biological information, IEEE/ACM transactions on computational biology and bioinformatics.

Zeng, M., et al. (2019) Protein-protein interaction site prediction through combining local and global features with deep neural networks, Bioinformatics.

Zhang, F., et al. (2019) DeepFunc: A Deep Learning Framework for Accurate Prediction of Protein Functions from Protein Sequences and Interactions, Proteomics, 1900019.

Zhang, T., et al. (2016) RNALocate: a resource for RNA subcellular localizations, Nucleic acids research, 45, D135–D138.

