## Supplemental tables and figures for "DeepLncLoc: a deep learning framework for long non-coding RNA subcellular localization prediction based on subsequence embedding"

---

**This supplementary file includes:**

**1. Supplementary Tables**

**Table S1.** Performance of different hyper-parameters (number of subsequences, k, dimension) in DeepLncLoc for prediction

**Table S2.** Distribution of the independent test dataset

**Table S3.** The detailed prediction results of DeepLncLoc, lncLocator, and iLoc-lncRNA on the test set

**Table S4.** Sequence length distribution of lncRNAs

**Table S5.** Species distribution of lncRNAs

### 2. Supplementary Figures

**Fig. S1.** Illustration of the five subcellular localizations included in the constructed dataset.

**Fig. S2.** Confusion matrices of DeepLncLoc with lncLocator on the test set. (a) lncLocator, (b) DeepLncLoc. Each row represents the true class while each column represents the predicted class.

**Fig. S3.** Confusion matrices of DeepLncLoc with iLoc-lncRNA on the test set. (a) iLoc-lncRNA, (b) DeepLncLoc. Each row represents the true class while each column represents the predicted class.

**Fig. S4.** The ACC and AUC of different species for DeepLncLoc

### 1. Supplementary Tables

**Table S1.** Performance of different hyper-parameters (number of subsequences, k, dimension) in DeepLncLoc for prediction

| No. subsequences | k | dimension | ACC | Macro F-measure | AUC |
|---|---|---|---|---|---|
| 16 | 1 | 64 | 0.511 | 0.337 | 0.802 |
| 16 | 1 | 128 | 0.512 | 0.333 | 0.801 |
| 16 | 2 | 64 | 0.548 | 0.417 | 0.817 |
| 16 | 2 | 128 | 0.562 | 0.404 | 0.821 |
| 16 | 3 | 64 | 0.547 | 0.411 | 0.819 |
| 16 | 3 | 128 | 0.562 | 0.414 | 0.821 |
| 16 | 4 | 64 | 0.551 | 0.377 | 0.815 |
| 16 | 4 | 128 | 0.557 | 0.402 | 0.822 |
| 16 | 5 | 64 | 0.544 | 0.387 | 0.819 |
| 16 | 5 | 128 | 0.551 | 0.375 | 0.814 |
| 16 | 6 | 64 | 0.518 | 0.353 | 0.797 |
| 16 | 6 | 128 | 0.531 | 0.376 | 0.799 |
| 32 | 1 | 64 | 0.519 | 0.366 | 0.805 |
| 32 | 1 | 128 | 0.526 | 0.375 | 0.804 |
| 32 | 2 | 64 | 0.560 | 0.413 | 0.817 |
| 32 | 2 | 128 | 0.546 | 0.395 | 0.813 |
| 32 | 3 | 64 | 0.541 | 0.393 | 0.815 |
| 32 | 3 | 128 | 0.562 | 0.416 | 0.816 |
| 32 | 4 | 64 | 0.551 | 0.404 | 0.819 |
| 32 | 4 | 128 | 0.537 | 0.381 | 0.816 |
| 32 | 5 | 64 | 0.545 | 0.389 | 0.806 |
| 32 | 5 | 128 | 0.552 | 0.367 | 0.810 |
| 32 | 6 | 64 | 0.538 | 0.362 | 0.809 |
| 32 | 6 | 128 | 0.536 | 0.362 | 0.809 |
| 64 | 1 | 64 | 0.533 | 0.385 | 0.812 |
| 64 | 1 | 128 | 0.520 | 0.387 | 0.796 |
| 64 | 2 | 64 | 0.547 | 0.409 | 0.813 |
| 64 | 2 | 128 | 0.548 | 0.403 | 0.814 |
| 64 | 3 | 64 | 0.548 | 0.421 | 0.819 |
| 64 | 3 | 128 | 0.549 | 0.412 | 0.818 |
| 64 | 4 | 64 | 0.545 | 0.384 | 0.805 |
| 64 | 4 | 128 | 0.551 | 0.418 | 0.819 |
| 64 | 5 | 64 | 0.531 | 0.368 | 0.802 |
| 64 | 5 | 128 | 0.540 | 0.367 | 0.807 |
| 64 | 6 | 64 | 0.533 | 0.374 | 0.808 |

| | | | | | |
|---|---|---|---|---|---|
| 64 | 6 | 128 | 0.533 | 0.366 | 0.813 |
| 128 | 1 | 64 | 0.518 | 0.381 | 0.802 |
| 128 | 1 | 128 | 0.517 | 0.387 | 0.806 |
| 128 | 2 | 64 | 0.559 | 0.433 | 0.812 |
| 128 | 2 | 128 | 0.547 | 0.422 | 0.818 |
| 128 | 3 | 64 | 0.537 | 0.387 | 0.819 |
| 128 | 3 | 128 | 0.567 | 0.432 | 0.821 |
| 128 | 4 | 64 | 0.539 | 0.400 | 0.813 |
| 128 | 4 | 128 | 0.553 | 0.410 | 0.824 |
| 128 | 5 | 64 | 0.526 | 0.384 | 0.818 |
| 128 | 5 | 128 | 0.551 | 0.400 | 0.817 |
| 128 | 6 | 64 | 0.527 | 0.368 | 0.814 |
| 128 | 6 | 128 | 0.527 | 0.362 | 0.814 |
| 256 | 1 | 64 | 0.529 | 0.393 | 0.812 |
| 256 | 1 | 128 | 0.507 | 0.392 | 0.799 |
| 256 | 2 | 64 | 0.537 | 0.404 | 0.818 |
| 256 | 2 | 128 | 0.534 | 0.423 | 0.807 |
| 256 | 3 | 64 | 0.544 | 0.388 | 0.824 |
| 256 | 3 | 128 | 0.546 | 0.407 | 0.817 |
| 256 | 4 | 64 | 0.538 | 0.402 | 0.818 |
| 256 | 4 | 128 | 0.546 | 0.395 | 0.823 |
| 256 | 5 | 64 | 0.548 | 0.400 | 0.819 |
| 256 | 5 | 128 | 0.535 | 0.347 | 0.814 |
| 256 | 6 | 64 | 0.520 | 0.359 | 0.810 |
| 256 | 6 | 128 | 0.552 | 0.354 | 0.817 |

**Table S2.** Distribution of the independent test dataset

| Subcellular localization | No. samples |
|---|---|
| Cytoplasm | 20 |
| Nucleus | 20 |
| Ribosome | 10 |
| Cytosol | 10 |
| Exosome | 7 |

**Table S3.** The detailed prediction results of DeepLncLoc, lncLocator, and iLoc-lncRNA on the test set

| No. | lncRNA | lncLocator | iLoc-lncRNA | DeepLncLoc | Label |
|---|---|---|---|---|---|
| 1 | >db_t_id=10162 \|chr12\|112277571\|112279706-\|ensembl_75_GRCh37\|Human\|ucsc_GCRh37\|Cytoplasm | Cytoplasm | Ribosome | Nucleus | Cytoplasm |
| 2 | >db_t_id=10210 \|chr2\|27293340\|27294517-\|ensembl_75_GRCh37\|Human\|ucsc_GCRh37\|Cytoplasm | Cytoplasm | Cytoplasm, Cytosol | Nucleus | Cytoplasm |
| 3 | >db_t_id=10265 \|chr1\|3652548\|3663886-\|ensembl_75_GRCh37\|Human\|ucsc_GCRh37\|Cytoplasm | Cytosol | Cytoplasm, Cytosol | Nucleus | Cytoplasm |
| 4 | >db_t_id=1028 \|chr10\|94196728\|94198949-\|ucsc_GCRm38\|Mouse\|ucsc_GCRm38\|Cytoplasm | Cytoplasm | Cytoplasm, Cytosol | Cytoplasm | Cytoplasm |
| 5 | >db_t_id=1029 \|chr5\|115358699\|115360123-\|ucsc_GCRm38\|Mouse\|ucsc_GCRm38\|Cytoplasm | Nucleus | Cytoplasm, Cytosol | Cytoplasm | Cytoplasm |
| 6 | >db_t_id=1032 \|chr9\|25136265\|25138587-\|ucsc_GCRm38\|Mouse\|ucsc_GCRm38\|Cytoplasm | Cytoplasm | Cytoplasm, Cytosol | Cytoplasm | Cytoplasm |
| 7 | >db_t_id=10403 \|chr6\|32862500\|32871535+\|ensembl_65_GRCh37\|Human\|ucsc_GCRh37\|Cytoplasm | Cytoplasm | Cytoplasm, Cytosol | Nucleus | Cytoplasm |
| 8 | >db_t_id=10666 \|chr1\|175126123\|175135877-\|ensembl_72_GRCh37\|Human\|ucsc_GCRh37\|Cytoplasm | Cytoplasm | Cytoplasm, Cytosol | Nucleus | Cytoplasm |
| 9 | >db_t_id=113 \|chr2\|113969099\|114034158+\|ensembl_75_GRCh37\|Human\|ucsc_GCRh37\|Cytoplasm | Cytoplasm | Cytoplasm, Cytosol | Nucleus | Cytoplasm |
| 10 | >db_t_id=11637 \|chr1\|20669882\|20684298+\|ensembl_91_GRCm38\|Mouse\|ucsc_GCRm38\|Cytoplasm | Cytoplasm | Cytoplasm, Cytosol | Cytoplasm | Cytoplasm |
| 11 | >db_t_id=1824 \|chr17\|62751554\|62752250-\|ucsc_GCRm38\|Mouse\|ucsc_GCRm38\|Cytoplasm | Cytoplasm | Cytoplasm, Cytosol | Cytoplasm | Cytoplasm |
| 12 | >db_t_id=8900 \|chr2\|90908716\|90918258-\|ensembl_91_GRCm38\|Mouse\|ucsc_GCRm38\|Cytoplasm | Cytoplasm | Cytoplasm, Cytosol | Nucleus | Cytoplasm |
| 13 | >db_t_id=9629 \|chr3L\|12627934\|12648280+\|ensembl_91_BDGP6\|FruitFly\|ucsc_BDGP6\|Cytoplasm | Nucleus | Nucleolus, Nucleus, Nucleoplasm | Nucleus | Cytoplasm |
| 14 | >db_t_id=9059 \|chr17\|29283298\|29283979-\|ensembl_75_GRCh37\|Human\|ucsc_GCRh37\|Cytoplasm | Cytoplasm | Nucleolus, Nucleus, Nucleoplasm | Cytoplasm | Cytoplasm |
| 15 | >db_t_id=9142 \|chr19\|48758932\|48761452+\|ensembl_65_GRCh37\|Human\|ucsc_GCRh37\|Cytoplasm | Cytoplasm | Cytoplasm, Cytosol | Nucleus | Cytoplasm |
| 16 | >db_t_id=9150 \|chr4\|103749212\|103765232+\|ensembl_75_GRCh37\|Human\|ucsc_GCRh37\|Cytoplasm | Cytoplasm | Cytoplasm, Cytosol | Nucleus | Cytoplasm |
| 17 | >db_t_id=9158 \|chr4\|139694704\|139722793+\|ensembl_75_GRCh37\|Human\|ucsc_GCRh37\|Cytoplasm | Cytoplasm | Cytoplasm, Cytosol | Nucleus | Cytoplasm |
| 18 | >db_t_id=9625 \|chr2L\|1380083\|1381326-\|ensembl_91_BDGP6\|FruitFly\|ucsc_BDGP6\|Cytoplasm | Nucleus | Cytoplasm, Cytosol | Cytoplasm | Cytoplasm |
| 19 | >db_t_id=9910 | Cytosol | Nucleolus, | Nucleus | Cytoplasm |

| | | | | | |
|---|---|---|---|---|---|
| | \|chr7\|100951627\|100954266+\|ensembl_75_GRCh37\|Human\|ucsc_GCRh37\|Cytoplasm | | Nucleus, Nucleoplasm | | |
| 20 | >db_t_id=9960 \|chr1\|235092978\|235095736+\|ensembl_75_GRCh37\|Human\|ucsc_GCRh37\|Cytoplasm | Cytoplasm | Cytoplasm, Cytosol | Nucleus | Cytoplasm |
| 21 | >db_t_id=10135 \|chr2\|74212259\|74213470+\|ensembl_75_GRCh37\|Human\|ucsc_GCRh37\|Nucleus | Exosome | Exosome | Ribosome | Nucleus |
| 22 | >db_t_id=10201 \|chr2\|27579113\|27583123+\|ensembl_75_GRCh37\|Human\|ucsc_GCRh37\|Nucleus | Cytosol | Cytoplasm, Cytosol | Nucleus | Nucleus |
| 23 | >db_t_id=1022 \|chr8\|25744389\|25745580-\|ucsc_GCRm38\|Mouse\|ucsc_GCRm38\|Nucleus | Cytoplasm | Nucleolus, Nucleus, Nucleoplasm | Cytoplasm | Nucleus |
| 24 | >db_t_id=10282 \|chr20\|37075221\|37079564+\|ensembl_75_GRCh37\|Human\|ucsc_GCRh37\|Nucleus | Cytoplasm | Cytoplasm, Cytosol | Nucleus | Nucleus |
| 25 | >db_t_id=10324 \|chr21\|26934221\|26947480+\|ensembl_75_GRCh37\|Human\|ucsc_GCRh37\|Nucleus | Cytoplasm | Cytoplasm, Cytosol | Nucleus | Nucleus |
| 26 | >db_t_id=10327 \|chr10\|75012549\|75014107+\|ensembl_75_GRCh37\|Human\|ucsc_GCRh37\|Nucleus | Nucleus | Nucleolus, Nucleus, Nucleoplasm | Nucleus | Nucleus |
| 27 | >db_t_id=10474 \|chr5\|33424131\|33440725-\|ensembl_75_GRCh37\|Human\|ucsc_GCRh37\|Nucleus | Cytoplasm | Nucleolus, Nucleus, Nucleoplasm | Nucleus | Nucleus |
| 28 | >db_t_id=10592 \|chr8\|144624143\|144631899-\|ensembl_75_GRCh37\|Human\|ucsc_GCRh37\|Nucleus | Ribosome | Ribosome | Nucleus | Nucleus |
| 29 | >db_t_id=11 \|chr4\|119199864\|119200978+\|ensembl_75_GRCh37\|Human\|ucsc_GCRh37\|Nucleus | Nucleus | Nucleolus, Nucleus, Nucleoplasm | Nucleus | Nucleus |
| 30 | >db_t_id=11029 \|chr6\|168224556\|168227389-\|ensembl_75_GRCh37\|Human\|ucsc_GCRh37\|Nucleus | Nucleus | Nucleolus, Nucleus, Nucleoplasm | Nucleus | Nucleus |
| 31 | >db_t_id=18 \|chr17\|18966762\|18967449-\|ensembl_75_GRCh37\|Human\|ucsc_GCRh37\|Nucleus | Cytoplasm | Ribosome | Ribosome | Nucleus |
| 32 | >db_t_id=2 \|chr6\|90539613\|90584155+\|ensembl_75_GRCh37\|Human\|ucsc_GCRh37\|Nucleus | Exosome | Exosome | Nucleus | Nucleus |
| 33 | >db_t_id=26 \|chr7\|45022622\|45026560-\|ensembl_75_GRCh37\|Human\|ucsc_GCRh37\|Nucleus | Nucleus | Cytoplasm, Cytosol | Nucleus | Nucleus |
| 34 | >db_t_id=4 \|chr17\|16342136\|16381992+\|ensembl_75_GRCh37\|Human\|ucsc_GCRh37\|Nucleus | Cytosol | Cytoplasm, Cytosol | Nucleus | Nucleus |
| 35 | >db_t_id=8840 \|chrX\|103460375\|103483217-\|ensembl_91_GRCm38\|Mouse\|ucsc_GCRm38\|Nucleus | Cytoplasm | Nucleolus, Nucleus, Nucleoplasm | Nucleus | Nucleus |
| 36 | >db_t_id=8933 \|chrX\|3847910\|3855896-\|ensembl_75_GRCh37\|Human\|ucsc_GCRh37\|Nucleus | Cytoplasm | Cytoplasm, Cytosol | Nucleus | Nucleus |
| 37 | >db_t_id=9009 \|chr1\|700237\|714006-\|ensembl_75_GRCh37\|Human\|ucsc_GCRh37\|Nucleus | Ribosome | Cytoplasm, Cytosol | Nucleus | Nucleus |
| 38 | >db_t_id=9146 \|chr14\|95998634\|96001137-\|ensembl_75_GRCh37\|Human\|ucsc_GCRh37\|Nucleus | Cytosol | Cytoplasm, Cytosol | Ribosome | Nucleus |
| 39 | >db_t_id=9698 | Cytoplasm | Cytoplasm, | Nucleus | Nucleus |

| | | | | | |
|---|---|---|---|---|---|
| | \|chr12\|109603940\|109661716+\|ensembl_91_GRCm38\|Mouse\|ucsc_GCRm38\|Nucleus | | Cytosol | | |
| 40 | >db_t_id=9710 \|chr11\|65265233\|65273940+\|ensembl_75_GRCh37\|Human\|ucsc_GCRh37\|Nucleus | Cytoplasm | Nucleolus, Nucleus, Nucleoplasm | Nucleus | Nucleus |
| 41 | >db_t_id=102 \|chr5\|175570088\|175626298-\|ensembl_75_GRCh37\|Human\|ucsc_GCRh37\|Ribosome | Cytoplasm | Cytoplasm, Cytosol | Nucleus | Ribosome |
| 42 | >db_t_id=119 \|chr4\|165675216\|165724947+\|ensembl_75_GRCh37\|Human\|ucsc_GCRh37\|Ribosome | Cytosol | Cytoplasm, Cytosol | Nucleus | Ribosome |
| 43 | >db_t_id=70 \|chr16\|2014960\|2015510+\|ensembl_75_GRCh37\|Human\|ucsc_GCRh37\|Ribosome | Cytosol | Cytoplasm, Cytosol | Nucleus | Ribosome |
| 44 | >db_t_id=87 \|chr2\|162101250\|162105561+\|ensembl_75_GRCh37\|Human\|ucsc_GCRh37\|Ribosome | Ribosome | Ribosome | Ribosome | Ribosome |
| 45 | >db_t_id=8935 \|chrX\|155244288\|155246502-\|ensembl_75_GRCh37\|Human\|ucsc_GCRh37\|Ribosome | Ribosome | Ribosome | Ribosome | Ribosome |
| 46 | >db_t_id=8942 \|chr20\|61405480\|61406764-\|ensembl_75_GRCh37\|Human\|ucsc_GCRh37\|Ribosome | Cytosol | Cytoplasm, Cytosol | Nucleus | Ribosome |
| 47 | >db_t_id=9029 \|chr11\|64413871\|64427153+\|ensembl_75_GRCh37\|Human\|ucsc_GCRh37\|Ribosome | Cytoplasm | Cytoplasm, Cytosol | Nucleus | Ribosome |
| 48 | >db_t_id=9047 \|chr9\|43102670\|43113375+\|ensembl_75_GRCh37\|Human\|ucsc_GCRh37\|Ribosome | Cytoplasm | Cytoplasm, Cytosol | Nucleus | Ribosome |
| 49 | >db_t_id=9188 \|chr5\|180688213\|180691262+\|ensembl_75_GRCh37\|Human\|ucsc_GCRh37\|Ribosome | Cytosol | Ribosome | Ribosome | Ribosome |
| 50 | >ID\|30352\|gene_id\|ENSG00000232019\|transcript_id\|ENST00000439105 AC074183.4 Homo sapiens lncRNA\|Ribosome | Nucleus | Nucleolus, Nucleus, Nucleoplasm | Ribosome | Ribosome |
| 51 | >db_t_id=30 \|chr5\|180618046\|180618852-\|ensembl_75_GRCh37\|Human\|ucsc_GCRh37\| Cytosol | Cytoplasm | Cytoplasm, Cytosol | Cytosol | Cytosol |
| 52 | >db_t_id=31 \|chr8\|67024902\|67027440+\|ensembl_75_GRCh37\|Human\|ucsc_GCRh37\| Cytosol | Cytosol | Cytoplasm, Cytosol | Cytosol | Cytosol |
| 53 | >db_t_id=32 \|chr8\|20133284\|20147969+\|ensembl_75_GRCh37\|Human\|ucsc_GCRh37\| Cytosol | Cytoplasm | Nucleolus, Nucleus, Nucleoplasm | Nucleus | Cytosol |
| 54 | >db_t_id=34 \|chr14\|23398818\|23424771+\|ensembl_75_GRCh37\|Human\|ucsc_GCRh37\| Cytosol | Ribosome | Ribosome | Nucleus | Cytosol |
| 55 | >db_t_id=27 \|chr17\|41447213\|41466567-\|ensembl_75_GRCh37\|Human\|ucsc_GCRh37\|Cytosol | Ribosome | Cytoplasm, Cytosol | Nucleus | Cytosol |
| 56 | >db_t_id=33 \|chr8\|144624280\|144624570+\|ensembl_75_GRCh37\|Human\|ucsc_GCRh37\|Cytosol | Cytosol | Cytoplasm, Cytosol | Cytosol | Cytosol |
| 57 | >db_t_id=45 \|chr22\|43011250\|43011913+\|ensembl_75_GRCh37\|Human\|ucsc_GCRh37\| Cytosol | Nucleus | Nucleolus, Nucleus, Nucleoplasm | Nucleus | Cytosol |
| 58 | >db_t_id=61 \|chr1\|17197440\|17200587+\|ensembl_75_GRCh37\|Human\|ucsc_GCRh37\| Cytosol | Cytosol | Cytoplasm, Cytosol | Cytosol | Cytosol |
| 59 | >db_t_id=72 \|chr16\|3206737\|3207484-\|ensembl_75_GRCh37\|Human\|ucsc_GCRh37\| Cytosol | Cytosol | Ribosome | Cytosol | Cytosol |
| 60 | >db_t_id=44 \|chr17\|62223330\|62223836+\|ensembl_75_GRCh37\|Human\|ucsc_GCRh37\|Cytosol | Exosome | Nucleolus, Nucleus, | Nucleus | Cytosol |

<table>
<tr><td></td><td></td><td></td><td>Nucleoplasm</td><td></td><td></td></tr>
<tr><td>61</td><td>>ID|14590|gene_id|407975|transcript_id|NR_027349 MIR17HG Homo sapiens lncRNA| Exosome</td><td>Nucleus</td><td>Cytoplasm,<br>Cytosol</td><td>Ribosome</td><td>Exosome</td></tr>
<tr><td>62</td><td>>ID|20872|gene_id|6023|transcript_id|NR_003051 RMRP Homo sapiens lncRNA| Exosome</td><td>Cytoplasm</td><td>Exosome</td><td>Exosome</td><td>Exosome</td></tr>
<tr><td>63</td><td>>ID|20873|gene_id|6029|transcript_id|NR_002715 RN7SL1 Homo sapiens lncRNA| Exosome</td><td>Cytoplasm</td><td>Exosome</td><td>Exosome</td><td>Exosome</td></tr>
<tr><td>64</td><td>>ID|2381|gene_id|112597|transcript_id|NR_024204 LINC00152 Homo sapiens lncRNA| Exosome</td><td>Cytosol</td><td>Ribosome</td><td>Cytosol</td><td>Exosome</td></tr>
<tr><td>65</td><td>>ID|30423|gene_id|L36162|transcript_id|L36162 L36162 Homo sapiens lncRNA| Exosome</td><td>Nucleus</td><td>Exosome</td><td>Exosome</td><td>Exosome</td></tr>
<tr><td>66</td><td>>ID|30602|gene_id|X15624|transcript_id|X15624 X15624 Homo sapiens lncRNA| Exosome</td><td>Nucleus</td><td>Cytoplasm,<br>Cytosol</td><td>Exosome</td><td>Exosome</td></tr>
<tr><td>67</td><td>>ID|8740|gene_id|23614|transcript_id|NR_002181 PPY2P Homo sapiens lncRNA| Exosome</td><td>Nucleus</td><td>Nucleolus,<br>Nucleus,<br>Nucleoplasm</td><td>Cytoplasm</td><td>Exosome</td></tr>
</table>

**Table S4.** Sequence length distribution of lncRNAs

| Sequence length | 0-1000 | 1000-3000 | 3000-7000 | 7000+ |
|---|---|---|---|---|
| Number | 177 | 384 | 157 | 139 |

**Table S5.** Species distribution of lncRNAs

| Species | Number |
|---|---|
| Mus musculus | 391 |
| Apis mellifera | 2 |
| Homo sapiens | 461 |
| Cricetulus griseus | 1 |
| Drosophila melanogaster | 1 |
| Gallus gallus | 1 |

### 2. Supplementary figures

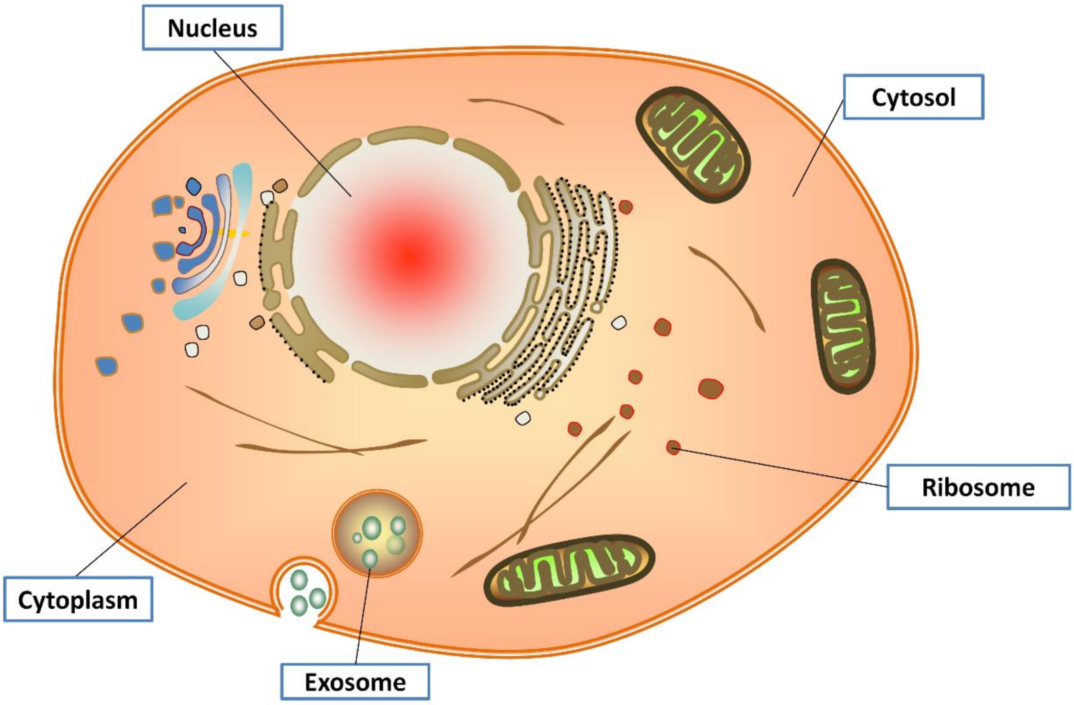


**Fig. S1.** Illustration of the five subcellular localizations included in the constructed dataset.

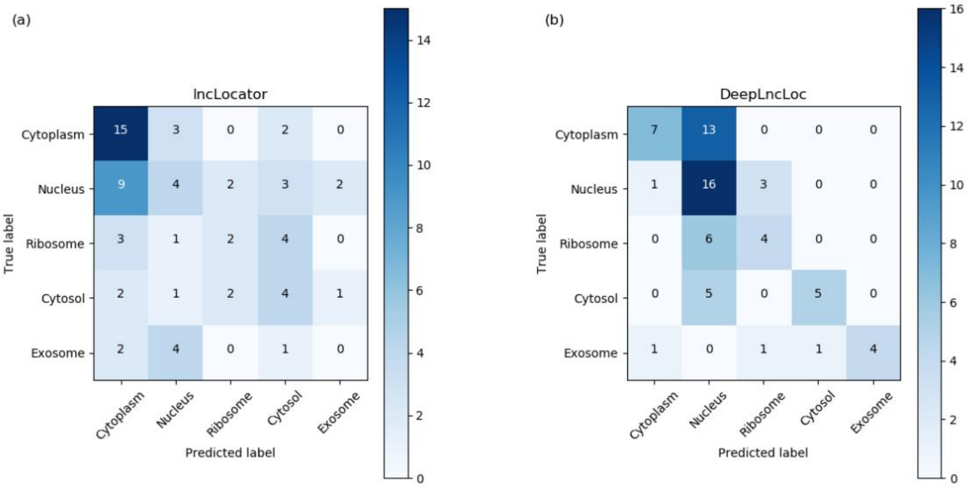


**Fig. S2.** Confusion matrices of DeepLncLoc with lncLocator on the test set. (a) lncLocator, (b) DeepLncLoc. Each row represents the true class while each column represents the predicted class.

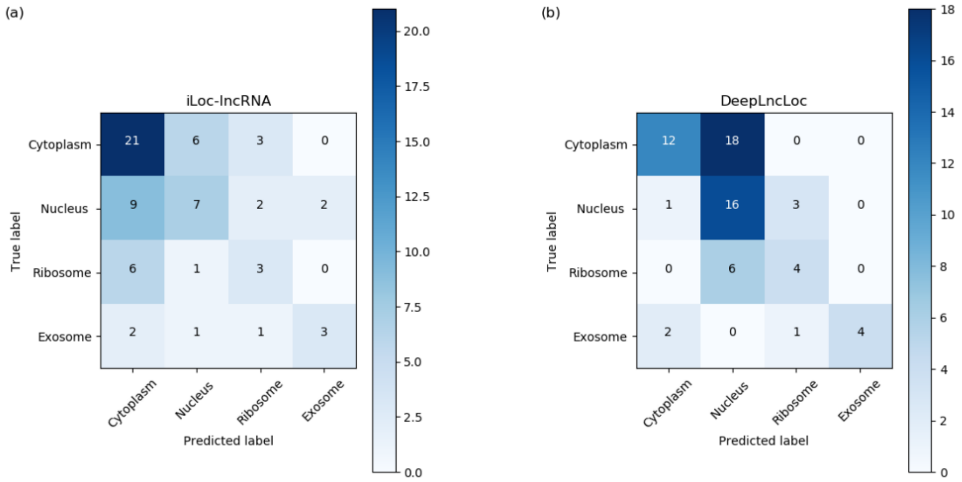


**Fig. S3.** Confusion matrices of DeepLncLoc with iLoc-lncRNA on the test set. (a) iLoc-lncRNA, (b) DeepLncLoc. Each row represents the true class while each column represents the predicted class.

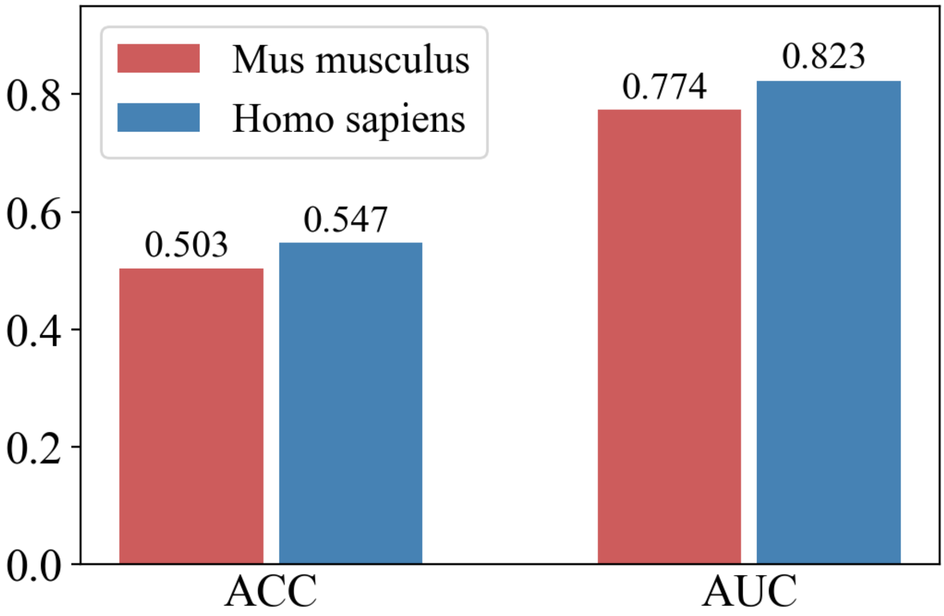


**Fig. S4.** The ACC and AUC of different species for DeepLncLoc
